# Odor Sampling Bags Enable Reliable Delivery of Controlled Odor Concentrations

**DOI:** 10.64898/2026.01.12.698699

**Authors:** Matthew Andres, Robert Pellegrino, Xuebo Song, Fernanda Ocampos, Joel D. Mainland

**Author notes:** These authors contributed equally (Matthew Andres, Robert Pellegrino).

## Abstract

Precise control of odorant concentration is essential for reliable olfactory research, yet existing odorant delivery methods often suffer from solvent interactions and dilution from ambient air, limiting stimulus consistency in olfactory research. We developed an odor sampling bag system using Nalophan plastic to create a closed headspace with air as the carrier medium, eliminating solvent-related variability and ambient air dilution. In two independent experiments, 15 trained panelists each rated the perceived intensity of seven concentrations of benzaldehyde and 2-heptanone using both gas-sampling bags and glass jars. Bags produced higher maximum perceptual intensities (p < 0.001) and greater test-retest reliability than jars (Experiment 1: r = 0.89 vs. 0.81, p < 0.001; Experiment 2: r = 0.86 vs. 0.72, p < 0.001). Notably, the two tested odorants showed different maximum intensities in bags (p < 0.001) but not jars (p = 0.85), suggesting bags better preserve odorant-specific concentration differences. Photoionization detector measurements confirmed stable headspace concentrations over time, comparable to industry-standard Tedlar bags. This cost-effective approach offers improved stimulus control for olfactory psychophysics research.

## Introduction

Careful control of sensory stimuli is essential for producing reliable and interpretable results in research. In olfaction, precise control of both odorant identity and concentration is necessary to accurately characterize the relationship between an odorant and its perception. However, inadequate stimulus control has led to inconsistencies in the literature: reported detection thresholds for the same odorant frequently differ by 4-5 orders of magnitude (Schmidt and Cain, 2010) and early researchers underestimated olfactory sensitivity due to variability in stimulus presentation (Gamble, 1898; Cain, 1977). Minimizing variability in olfactory stimuli improves the accuracy and reliability of psychophysical measurements.

Researchers have developed various odorant delivery methods, but most fail to provide absolute concentration control. Static methods—including sealed jars containing odorants diluted in solvent and impregnated pens (e.g. Sniffin’ Sticks) (Hummel et al., 1997)—lack standardization of delivery vehicles and solvents, preventing precise concentration control (Albrecht et al., 2008). Dynamic olfactometry uses devices to present odors in controlled airflow, enabling precise timing (Lundström et al., 2010) or control of the output concentration (Johnson and Sobel, 2007), but due to dilution from ambient air during a sniff most systems do not control the absolute concentration reaching the olfactory epithelium. Although these methods suit many applications, precise stimulus control becomes essential for rigorous investigation of concentration-perception relationships.

Three fundamental problems limit concentration control in existing methods: solvent interactions, incompatible solvent mixtures, and dilution with external air. The relationship between liquid-phase and vapor-phase concentration often deviates from theoretical predictions due to solvent interactions, with vapor concentration differing by at least an order of magnitude depending on whether water or mineral oil is used as solvent (Jennings et al., 2022). Studying odor mixtures introduces additional complications, as differing polarity among odorants limits solvent choices. Finally, sampling from an open system introduces external air, diluting vapor concentration and introducing ambient odors.

To address these issues, we developed a novel method using gas-sampling bags made of Nalophan plastic to deliver odors to human subjects. Our system uses air as a universal odorant solvent and eliminates the complications of liquid-to-gas-phase conversions. The closed system prevents introduction of external air during sampling, as the bag collapses with each sniff, avoiding dilution. Gas-sampling bags are standard tools in environmental odor analysis, and the Triangle Odor Bag method, which employs human panels to sample from bags for environmental odor assessment, was developed in Japan over 50 years ago (Iwasaki, 1972, 2012). Abraham and colleagues (Abraham et al., 2012) assumed that this approach provided superior concentration control in a comprehensive detection threshold study, as it reported lower detection thresholds, but there was no quantitative comparison to the more typical static olfactometry. Our system adapts bag-based delivery specifically for psychophysical research at lower cost than existing gas-sampling bags.

To validate this method, we compared perceptual intensity ratings between bags and traditional jars across two experiments with independent cohorts, hypothesizing that bags would produce higher maximum intensities and greater test-retest reliability. We also measured headspace concentrations using photoionization detection to objectively assess concentration stability over time.

## Materials and Methods

### Subjects

Two independent groups participated in Experiments 1 and 2, respectively. Experiment 1 included 15 individuals (11 females, mean age 34.5 ± 11.4 years), and Experiment 2 included 15 individuals (8 females, mean age 34.1 ± 11.5 years). All participants reported a healthy sense of smell, were 55 years of age or younger, and had no conditions associated with olfactory impairment (e.g., renal disease, neurodegenerative disease, or nasal congestion) at the time of testing.

### Odor Sampling Bag

The bag (Figure 1) is composed of Nalophan, a thin plastic material commonly used for gas sampling bags (Miller and McGinley, 2008). One end is sealed around a tube fitted with a septum for odorant injection; the other is sealed around an open/close valve for airflow control. Both ends are secured with cable ties. To prepare a bag, we first evacuated the air by attaching the valve to a vacuum line. We then filled the bag with dehumidified, carbon-filtered air at a known flow rate (e.g., 2 L/min) for a set duration (e.g., 5 min), yielding a known volume (e.g., 10 L). Liquid odorant was then injected through the septum via syringe. Injection volumes ranged from 0.5 µL to 2 mL depending on bag volume, target concentration, and the odorant’s volatility. For example, achieving a headspace concentration of 3 × 10^− 6^ (v/v) in a 10 L bag required injecting 30 µL of liquid odorant. Equilibration time varied from minutes to hours depending on the volume and volatility of the injected liquid.

**Figure 1.**
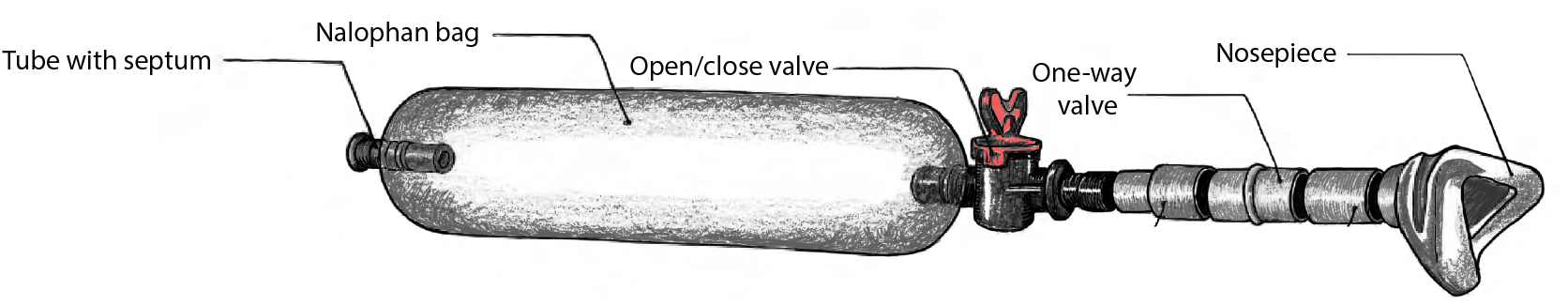
Odor sampling bag schematic. The bag is constructed from thin Nalophan tubing, sealed at one end with a septum for odorant injection and at the other with an open/close valve for attachment to a nose mask. The closed system maintains a consistent headspace concentration across repeated sampling without dilution.

For sampling, the valve was connected to a nose mask fitted with a low-resistance one-way valve, allowing participants to sniff naturally while preventing external air from entering the bag. The soft, flexible mask formed a seal against the face and could be quickly attached and detached, enabling a single mask to sample from multiple bags. A complete materials list and assembly instructions are provided in the supplementary material.

### Odor Stimuli

Two odorants were used, 2-heptanone and benzaldehyde, for their broad perceptual intensity ranges. For each odorant, seven concentrations were chosen based on preliminary data to span the concentration-intensity curve, with the two highest concentrations targeting perceptual saturation and the lowest targeting near-threshold detection.

Seven 10 L bags were prepared for each odorant (one per concentration) the day before testing began, plus one bag of non-odorized air as a control. For jar stimuli, seven concentrations of each odorant were prepared using propylene glycol as the diluent. In Experiment 1, 0.5 mL of each dilution was pipetted onto cellulose beads in 30 mL glass jars. In Experiment 2, 5 mL of each dilution was pipetted directly into 500 mL glass jars without beads. All procedures were conducted according to the Declaration of Helsinki for studies on human subjects and approved by the University of Pennsylvania IRB review for research involving human subjects (IRB #818208).

### Experiment 1

Experiment 1 compared perceptual intensity ratings between the bag system and traditional jar delivery. Fifteen participants rated the intensity of all seven concentrations of each odorant in both delivery formats using the generalized Labeled Magnitude Scale (gLMS) in duplicate. Prior to testing, participants were trained on the gLMS using examples from other sensory modalities and instructed to rate intensity independent of pleasantness. During the ∼1-hour session, participants sampled from both bags and jars, with presentation order counterbalanced across participants. A minimum 30-second interval separated each sample to minimize adaptation.

### Experiment 2

Experiment 2 replicated Experiment 1 with an independent cohort of 15 participants, using 500 mL jars instead of 30 mL jars. This modification tested whether any observed differences between bags and jars in Experiment 1 were attributable to limited headspace volume in the smaller jars. All stimuli were rated in duplicate.

### PID Measurements

Headspace concentrations were quantified using a photoionization detector (PID). The PID was calibrated daily with certified isobutylene gas to verify instrument accuracy and precision and to minimize measurement drift, according to the manufacturer’s recommendation. To convert PID voltage to parts per million (PPM), we generated a multi-point calibration curve by measuring known concentrations of each odorant via air dilution, with saturated headspace concentration calculated from vapor pressure. Headspace measurements were then taken from bags and jars at the same concentrations used in psychophysical testing, and PPM values were interpolated from the calibration curve.

To assess concentration stability of the Nalophan bags, we measured samples over a four-day period beginning one day after preparation. Measurements were taken under varying conditions: odorant (2-heptanone vs. benzaldehyde), concentration level (high vs. low), and bag type (Nalophan vs. Tedlar). Two bags of each type were prepared and measured.

PID accuracy was further assessed with a common gas standard (500 PPM of isobutylene) at 500 PPM, and diluted using mass flow controllers to 250 PPM, and 100 PPM. Three bags of each concentration were prepared and measured.

### Statistical Analysis

#### Reliability

Test-retest reliability was assessed for each experiment using two metrics: root mean squared error and Pearson correlation between duplicate ratings. Correlations between delivery types within each session were compared using Silver, Hittner, and May’s (Silver et al., 2004) modification of Dunn and Clark’s (Dunn and Clark, 1969) z-test with back-transformed averaged Fisher’s Z values (cocor package; (Diedenhofen and Musch, 2015)). Changes in correlation across sessions within a delivery type were compared using Fisher’s Z test.

#### Concentration-intensity function

Data were collapsed across experiment and grouped by delivery type and odorant, then fitted with a Hill function which accurately models human intensity ratings (Chastrette et al., 1998). Initial fits revealed poor model performance; therefore, a normalization and bootstrap procedure was completed. Intensity ratings were normalized to control for individual differences in scale usage using a geometric-mean rescaling procedure after Bartoshuk et al. (2004). Next, uncertainty in the Hill Equation parameters was assessed using nonparametric bootstrapping (N = 50 resamples), in which participants were resampled with replacement within each odor and delivery type while preserving within-subject observations. The Hill model was refit for each bootstrap sample, yielding distributions of parameter estimates and predicted responses that were used to estimate variability and confidence intervals. Differences in delivery type among hill parameters were assessed with a linear mixed model with parameters as the outcome, delivery type as the fixed effect, and odorant as a random effect (lme4 package, (Bates et al., 2015)).

#### Intensity comparisons

Differences in intensity ratings between bags and jars were tested using a linear mixed model with intensity as the outcome, the odorant × delivery type interaction as fixed effects, and participant as a random effect (lme4 package). Significant interactions were followed by simple-effects comparisons using estimated marginal means with Tukey adjustment for multiple comparisons (emmeans, (Lenth and Piaskowski, 2025)).

## Results

### Perceptual Intensity Ratings

Dose-response curves were constructed by plotting perceptual intensity ratings against PID-measured concentrations (PPM) for 2-heptanone and benzaldehyde (Figure 2). Curve shapes differed significantly between bags and jars. For both odorants, bags produced higher maximum perceptual intensities than jars at equivalent concentrations [bags (mean, sd): 42.5 (5.3) jars: 22.0 (2.8); F(197) = 2476.1, p < 0.001]. The two highest concentrations yielded significantly different intensity ratings between odorants for bags (F(436) = 4.2, p < 0.001) but not for jars [F(435) = 0.2, p = 0.85]. This result was expected, as the closed bag system prevents ambient air from diluting the sample during sniffing.

**Figure 2.**
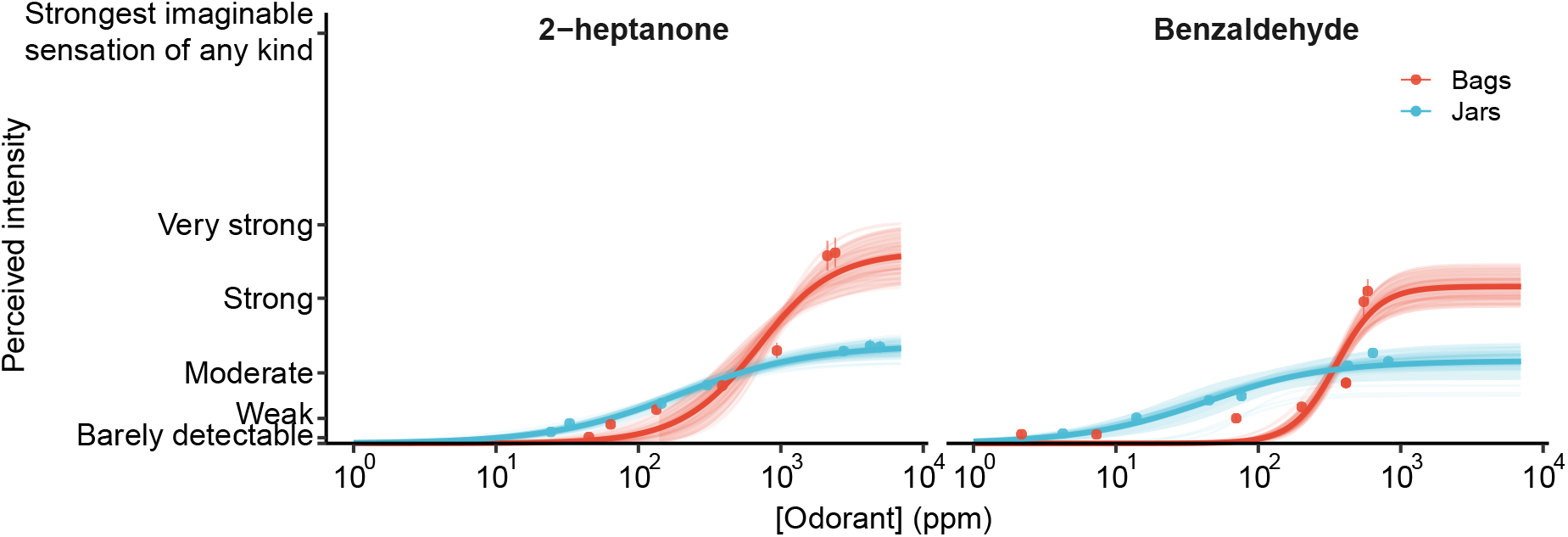
Concentration-intensity functions for bags and jars. Bag presentations (N = 30 subject–sessions; 50 bootstrap resamples) exhibited higher asymptotic intensity estimates and steeper slopes than jar presentations for both odorants (p < 0.001). Jar presentations reached perceptual saturation at lower intensities across concentrations. Estimated EC50 values also differed between delivery types (p < 0.001).

Unexpectedly, both odors had lower EC50s in jars compared to bags indicating earlier detection [bags (mean, sd): 512.4 (186.9) ppm, jars: 120.2 (81.8) ppm; F(197) = 1205.5, p < 0.001], but EC50 depends strongly on max intensity and slope. As shown the max intensity was greater for bags than jars. Similarly, bags exhibited steeper slopes than those obtained in jars [bags (mean, sd): 2.3 (.7), jars: 1.0 (.2); F(197) = 554.2, p < 0.001], indicating greater perceptual sensitivity to concentration changes and enhanced distinguishability across odor intensities.

### Reliability of Ratings

All stimuli were rated in duplicate, and within-participant consistency served as a measure of test-retest reliability. Both delivery methods produced reliable ratings across experiments; however, bags yielded significantly higher reliability than jars in both Experiment 1 (r = 0.89 vs. 0.81; z = 3.43, p < 0.001) and Experiment 2 (r = 0.86 vs. 0.72; z = 4.21, p < 0.001; Figure 3).

**Figure 3.**
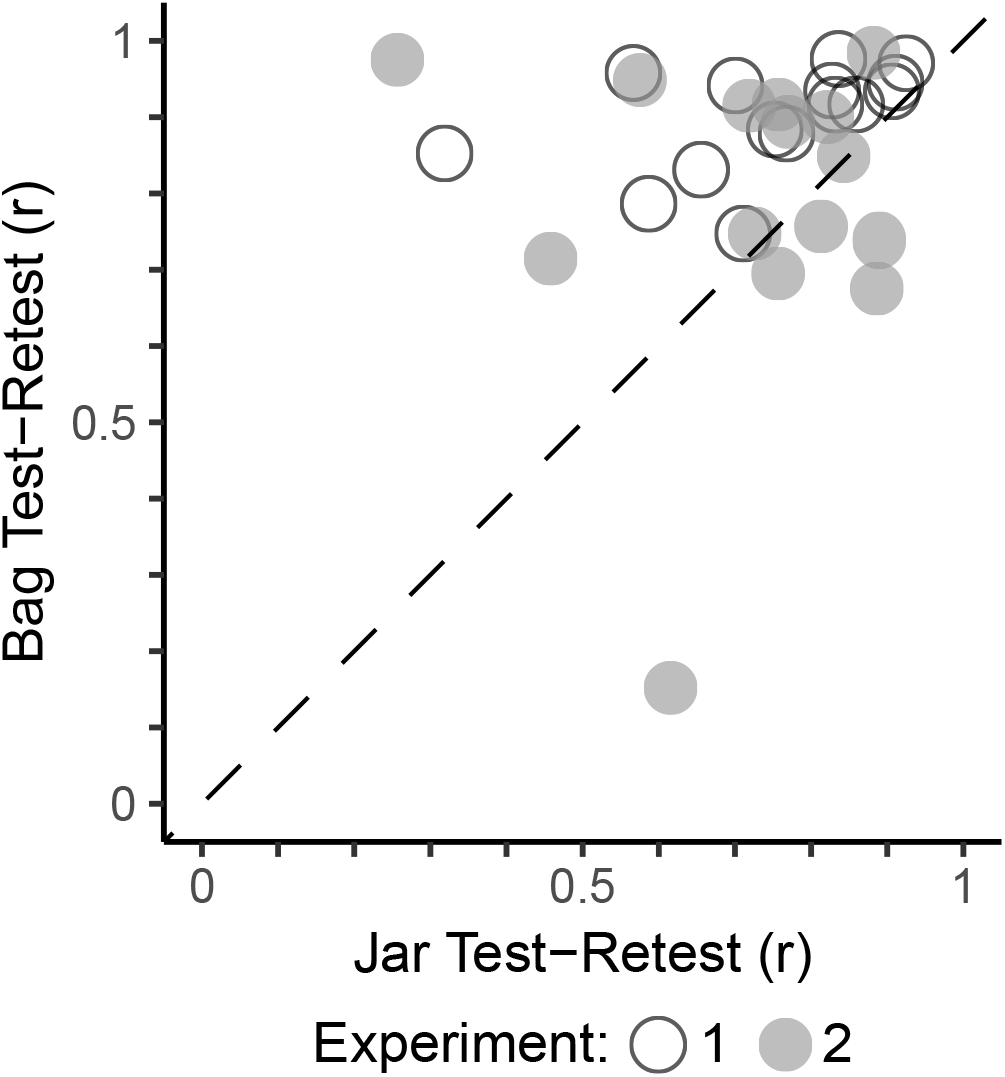
Test-retest reliability by delivery method. Participants showed higher within-session reliability for bags than jars in both Experiment 1 (open circles; p < 0.001) and Experiment 2 (close circles; p < 0.001).

Reliability remained stable across experiments for bags (z = 1.31, p = 0.19) but declined significantly for jars (z = 2.25, p = 0.03).

### Shelf-life of Bags

Nalophan bags maintained headspace concentrations over time at rates comparable to industry-standard Tedlar bags (Figure 4). When identical volumes of liquid odorant were injected, both bag types produced equivalent initial headspace concentrations. Lower concentrations remained stable across the four-day measurement period, whereas higher concentrations showed some loss. Concentration stability also varied by odorant, with 2-heptanone showing greater stability than benzaldehyde.

**Figure 4.**
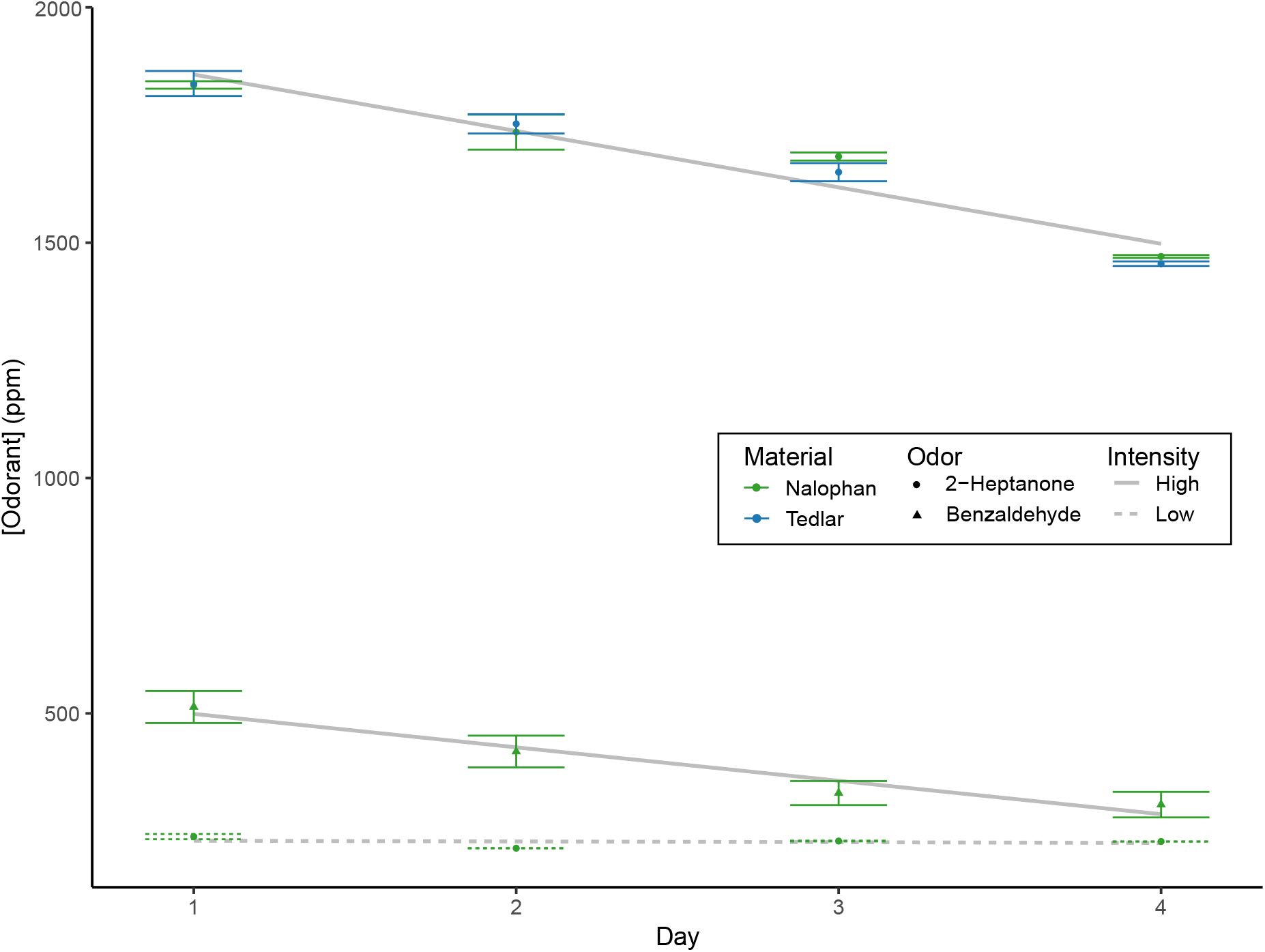
Headspace concentration stability over time. PID-measured concentrations for high concentration 2-heptanone in Nalophan, compared with substituting Tedlar bags (blue), benzaldehyde (triangles), and a low concentration (dotted line) across four days (in duplicate). Both bag types showed comparable stability, with greater loss at higher concentrations. Error bars represent standard error.

Measurement accuracy was assessed using isobutylene gas (Figure 5). Measurement variability was low across all concentrations. At lower and intermediate concentrations, measured values closely matched the nominal concentrations, showing minimal bias. At the highest concentration levels, a small but consistent negative deviation from the nominal concentration was observed, indicating a slight systematic bias at the upper end of the measurement range.

**Figure 5.**
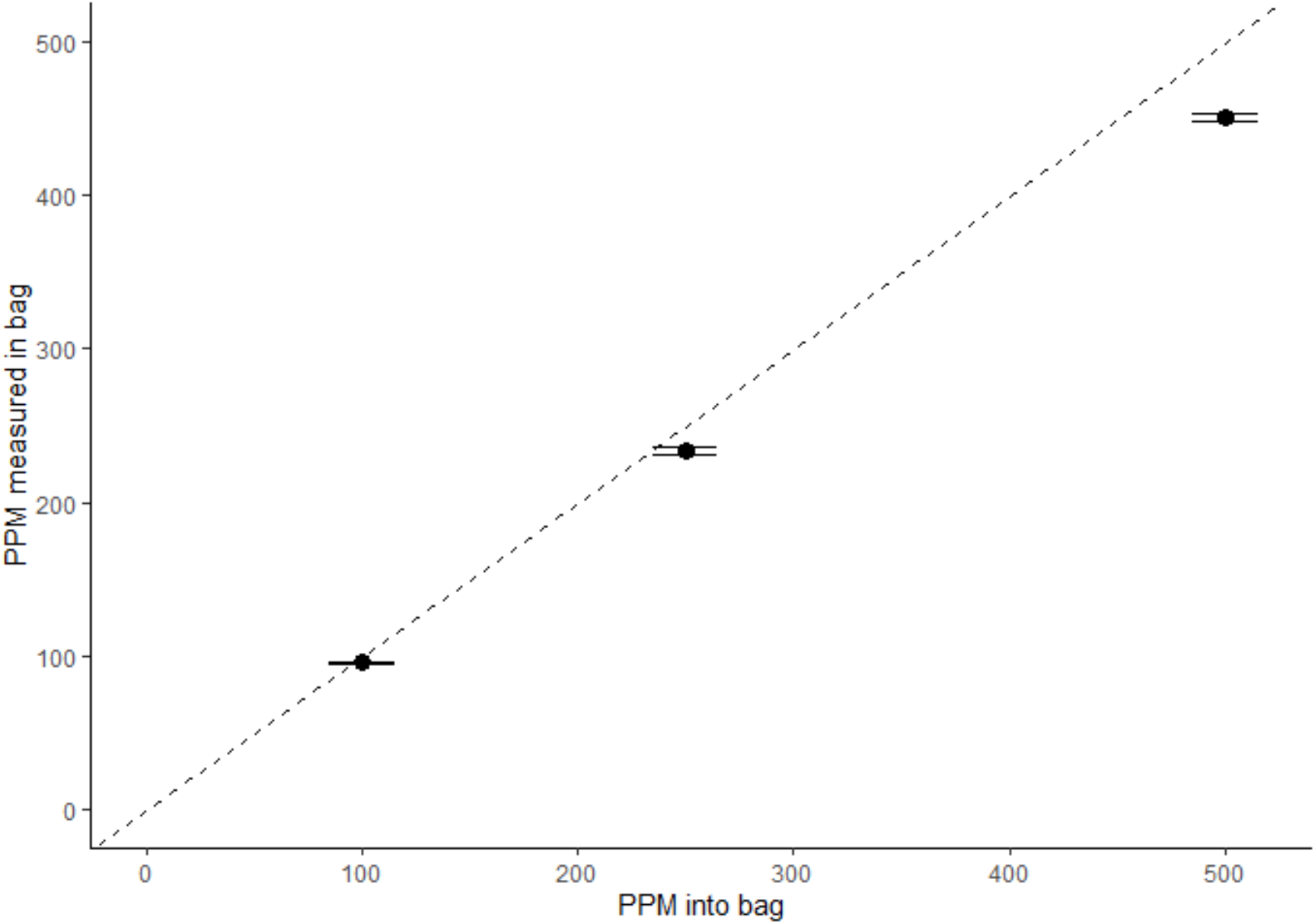
Measured versus expected headspace concentration. PID-measured three concentrations of isobutylene in bags (in triplicate) plotted against expected concentrations based on injected concentration (500, 250 and 100 ppm). Error bars represent standard error of bag measures.

## Discussion

We have demonstrated that our bag-based odorant delivery system produces more reliable and higher-intensity perceptual ratings than traditional jar methods. Consistent concentration across samples yielded high test-retest reliability, increasing confidence in psychophysical measurements. Bags may be particularly advantageous in studies with small sample sizes, where minimizing measurement noise is critical. We attribute this consistency to the elimination of variation in air dilution and ambient odorants during sampling. By bypassing liquid-phase dilution entirely, the bag system allows direct control of vapor concentration, making it well suited for experiments requiring precise stimulus control—including studies of odor mixtures, where solvent compatibility constraints often limit traditional approaches.

The bag system also achieved higher maximum perceptual intensities than jars at equivalent vapor concentrations, even when jar volume was increased to provide sufficient headspace. This makes bags particularly useful for studies requiring perception of very intense odors or for sampling typically weak odorants at their highest achievable intensity.

Odor concentration–intensity functions showed steeper slopes in bags than in jars, while mid-range intensities were higher in jars. Steeper slopes are consistent with greater sensitivity to concentration changes and increased perceptual discriminability. In contrast, estimated inflection points (EC50) suggest lower detection levels in bags, a pattern that differs from prior reports (Abraham et al., 2012) and may reflect the strong dependence of EC50 on slope and asymptotic intensity. Together, these results indicate presentation-dependent changes in odor intensity coding whose underlying mechanisms remain unresolved. Future studies examining a wider set of odorants spanning systematic differences in volatility, polarity, and related physicochemical properties may help elucidate the basis of these effects.

More broadly, this method has the potential to improve cross-laboratory replicability in olfactory research. The variability in stimulus control across current methods contributes to inconsistencies in the literature, such as detection thresholds for the same odorant differing by orders of magnitude. A standardized, low-cost delivery system with verified concentration stability could help address this problem, enabling more accurate threshold measurements and facilitating comparison of results across studies.

Several limitations of the bag method should be noted. Preparing samples is more time-consuming than liquid dilution, and bags have a finite shelf life due to cumulative odorant loss in the absence of a liquid reservoir. Nalophan absorbance can differ among odorants; therefore we recommend evaluating shelf life across experimental odorants. The method also requires manual administration by the experimenter, unlike self-administered jar testing or automated olfactometry. Finally, extending this approach to non-human animal subjects would require additional development. The present validation was also limited in scope: only two odorants were tested, and sample sizes were modest. Generalizability to other odorants and experimental paradigms remains to be established.

In summary, our bag system is an effective technique for collecting psychophysical data, particularly for studies dependent on sampling consistency, stimulus control, and concentration precision. While not suited to all applications, it offers a practical, cost-effective alternative to existing methods when precise control of air-phase concentration is paramount.

## Supporting information

Supplementary Protocol

## Acknowledgements

We would like to thank Adam Dewan for help with PID calibration techniques, Gregory Lane and Christina Zelano for help with component design and sourcing, Jennifer Margolis and Federica Genovese for helpful conversations and the George Preti Research Support Core for Analytical Chemistry for guidance.

