## Supplementary Protocol for "Odor Sampling Bags Enable Reliable Delivery of Controlled Odor Concentrations"

### Supplemental Protocol S1: Odorant Delivery Bag Assembly and Operation

#### Overview

This protocol describes the construction of sampling bags for controlled odorant delivery in psychophysical and neuroimaging experiments. The bags consist of Nalophan tubing sealed at both ends: one end fitted with an open/close valve for air flow and participant delivery, and the other end fitted with a septum port for odorant injection. Figure S1 illustrates the complete assembly and delivery system.

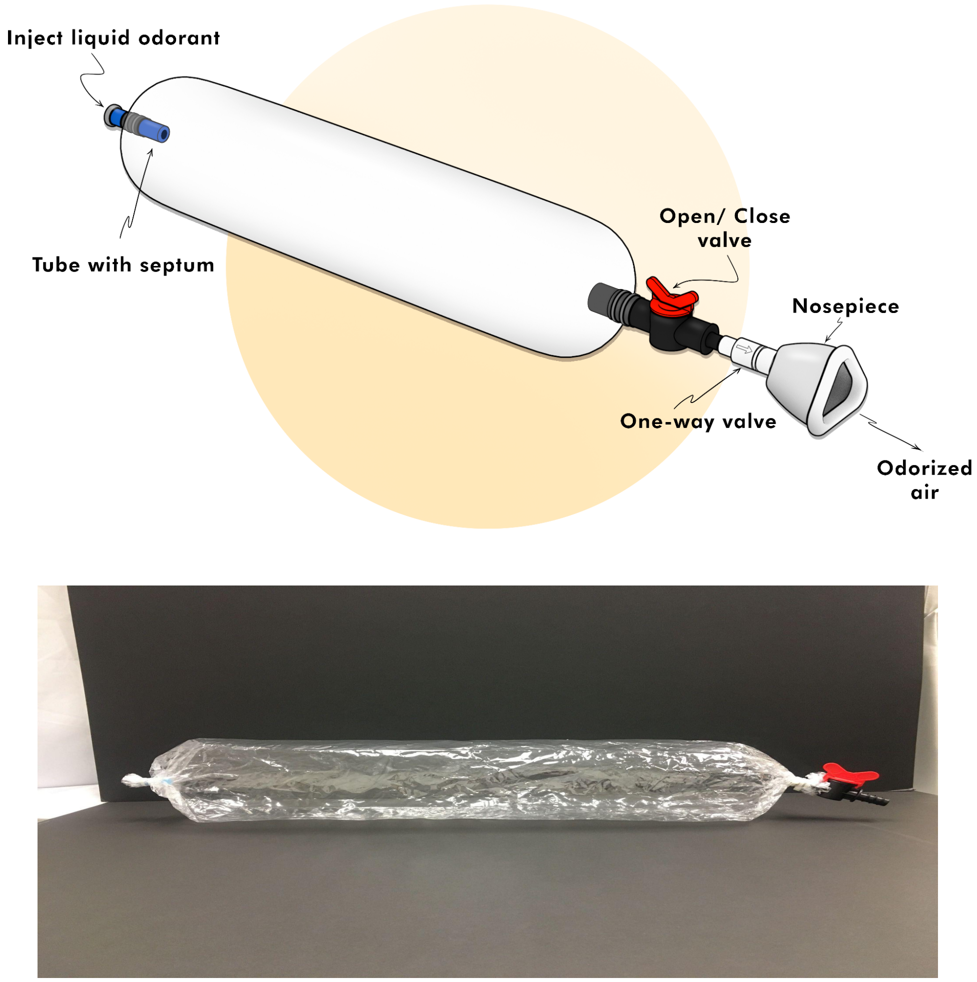


**Figure S1.** Schematic diagram (top) and photograph (bottom) of the complete odorant delivery bag system, showing the septum injection port, Nalophan bag, open/close valve, one-way valve, and nosepiece assembly.

#### Materials

##### Bag Components

| Component | Supplier | Catalog No. |
|---|---|---|
| Nalophan bags | Scentroid | — |
| Silicone tubing (5/16" ID, 7/16" OD) | McMaster-Carr | 9334T53 |
| Aluminum crimp caps with septum | Thomas Scientific | 2679H31 |
| Cable ties (clear) | Various suppliers | — |
| Open/close valve | U.S. Plastic Corp | 36488 |
| Parafilm M | Thomas Scientific | 21A00J411 |

##### Tools

| Tool | Supplier | Catalog No. |
|---|---|---|
| Cable tie gun | Uline | H-241 |
| Manual crimp cap crimper | Thomas Scientific | 1184L67 |
| Tubing cutter (optional) | Various suppliers | — |

##### Delivery System Components

| Component | Supplier | Catalog No. |
|---|---|---|
| Nasal mask | Amazon (Dreamwear) | B00JG1APG8 |
| Silicone tubing (7/8" × 15/16") | Cole-Parmer | EW-96202-93 |
| One-way valve | CareFusion AirLife (Medex Supply) | 104478 |
| Headgear frame | Amazon | B00D8BBFFW |
| Headgear straps | Amazon | B00CF4JJLK |

##### Bag Dimensions

Table S1 provides recommended Nalophan bag lengths for target volumes.

| Bag Length (cm) | Maximum Volume (L) |
|---|---|
| ~50 | 4.5 |
| ~70 | 8 |
| ~90 | 10 |
| ~130 | 20 |
| ~190 | 30 |

**Table S1.** Recommended Nalophan bag lengths for target volumes.

#### Bag Assembly Procedure

##### Valve End Assembly

1. **Cut the Nalophan bag to the desired length** (see Table S1 for volume estimates).
2. **Prepare the open/close valve.**
   a. Wrap at least one layer of Parafilm around one end of the valve to increase friction and improve the seal.
3. **Attach the bag to the valve.**
   a. Position the valve at one end of the bag opening, with the Parafilm-wrapped end inside.
   b. Fold the bag material over the valve tube so the opening rim is positioned near the valve's center piece (Figure S2).
   c. ***Critical:*** Ensure all folds are aligned; displaced folds will cause air leaks.

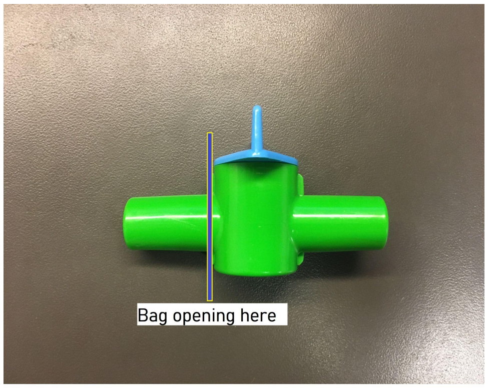


**Figure S2.** Correct positioning of bag material on the open/close valve, showing fold alignment near the valve center piece.

4. **Secure the bag with cable ties.**
   a. Fasten the first cable tie around the bag onto the middle of the valve tube. Use a cable tie gun to secure tightly and trim excess.
   b. Add a second cable tie approximately 6 mm (~¼ inch) closer to the bag opening, ensuring the material between ties is taut.
   c. Add a third cable tie 6 mm above the second.
   d. ***Critical:*** The bag opening rim must be entirely above the uppermost cable tie. If not, carefully cut the ties without puncturing the bag and restart.

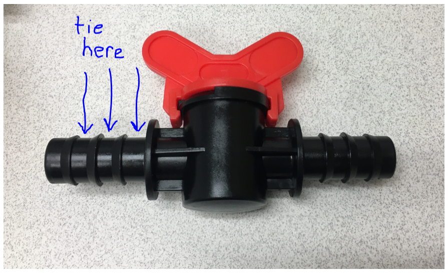


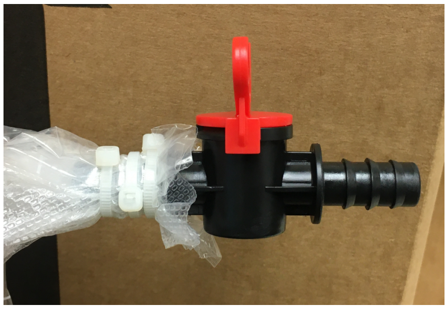

**Figure S3.** Cable tie placement on the valve end, showing three ties securing the bag to the valve tube.

**Septum End Assembly**

5. **Cut the silicone tubing.**
    a. Cut a 3.5–4 cm segment of silicone tubing (5/16" ID, 7/16" OD). Ensure the cut is straight and perpendicular. A tubing cutter produces the cleanest results but is not required.
6. **Attach the septum cap.**
    a. Place an aluminum crimp cap with septum on one end of the tubing segment.
    b. Use a manual crimping tool to crimp the cap onto the tube. The cap should form an airtight seal with no gaps at either the top or bottom edges (Figure S4).

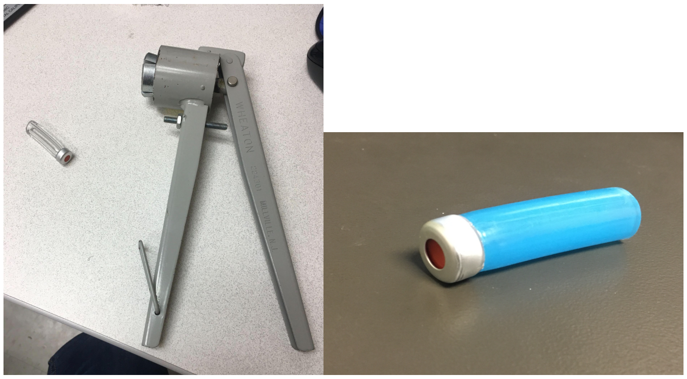

**Figure S4.** Septum cap crimping tool (left) and properly sealed septum tube (right).

7. **Attach the septum tube to the bag.**
    a. Attach the septum tube to the opposite end of the bag using the same cable tie method as the valve end.
    b. Place the first cable tie at the middle of the septum tube, then add one tie on each side.
    c. ***Critical:*** Ensure the tube opening remains unobstructed and no bag material covers the interior opening.
    d. Trim any excess bag material around the septum if necessary (Figure S5).

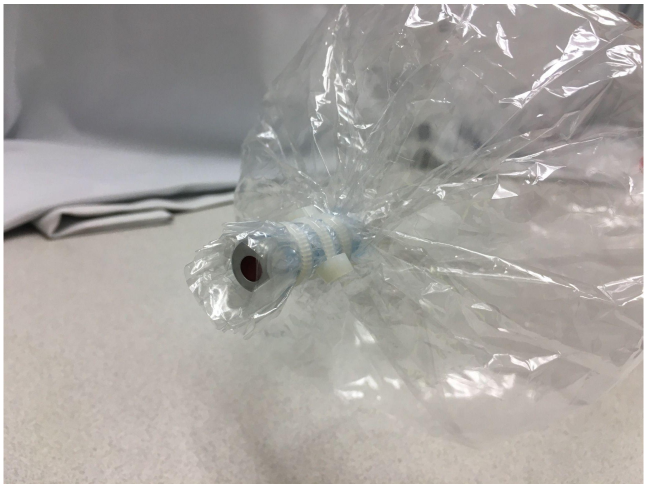

**Figure S5.** Completed septum end assembly showing proper cable tie placement and trimmed excess material.

#### Bag Filling Procedures

##### Liquid Odorants

###### *Air Evacuation*

1. Connect tubing to the open/close valve.
2. Open the valve and activate the vacuum source.
3. Once air is evacuated from bag folds, close the valve before disconnecting the vacuum.

###### *Air Filling*

4. Set the desired flow rate using an air flow regulator and flow meter or mass flow controller (e.g., 2 L/min).
5. Connect tubing to the open/close valve.
6. Open the valve and start a timer. Fill for the calculated duration based on target volume and flow rate (e.g., for 10 L at 2 L/min: 10 L ÷ 2 L/min = 5 min). We automate this process with a mass flow controller and custom software.
7. Close the valve and disconnect tubing.

###### *Odorant Injection*

8. Fill a syringe with the liquid odorant, ensuring no air bubbles are present. Note that odorant viscosity varies.
9. Insert the needle through the septum and inject the odorant into the bag.
   a. ***Caution:*** Avoid puncturing the bag material with the needle.
   b. Orient the bag to prevent liquid from pooling in folds and creases (Figure S6).

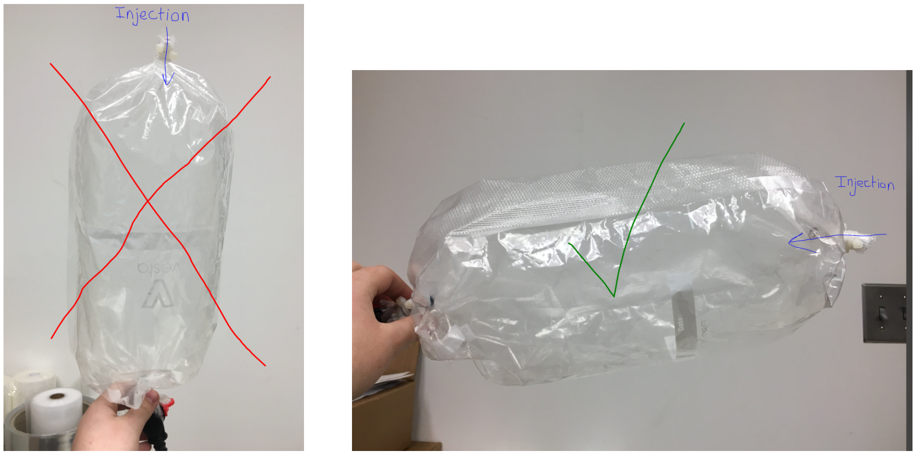


**Figure S6.** Bag orientation during odorant injection. Left: Incorrect orientation allowing liquid to pool in folds. Right: Correct orientation promoting even distribution.

10. Allow sufficient time for liquid evaporation (typically overnight, ranging from minutes to hours depending on the odorant).

    ***Note:*** We typically prepare bags the day before testing to ensure complete evaporation. Residual liquid visible on the bag surface indicates incomplete volatilization or excessive odorant quantity.

**Gaseous Odorants**

For odorants supplied as compressed gases, use the following procedure to prepare diluted mixtures:

1. Connect clean air and odorant gas tanks to the mixing regulator via PVC tubing.
2. Ensure regulator valves are closed (clockwise until resistance).
3. Fully open tank valves (counterclockwise) for both air and odorant gas. Verify equal pressure readings (typically 20 psi).
4. Connect the mixture outlet to a flow meter.
5. Calculate the required air flow rate: Air flow = (1 – target fraction) × total flow rate.

   *Example:* For 10% odorant at 2 L/min total: Air = (1 – 0.10) × 2 = 1.8 L/min
6. Slowly open the air valve to achieve the calculated air flow rate.
7. Slowly open the odorant gas valve until the total flow rate reaches the target value.
8. Disconnect from the flow meter and connect to the evacuated bag.
9. Open the bag valve and fill to the desired volume (e.g., 2 L/min for 1 min = 2 L).
10. Close the bag valve, then close the regulator valves.
11. Close tank valves if not in continued use.

#### Odorant Delivery System

The delivery system connects the filled odorant bag to a nasal mask worn by the participant (Figure S7). This configuration ensures controlled, reproducible odorant presentation during experimental sessions.

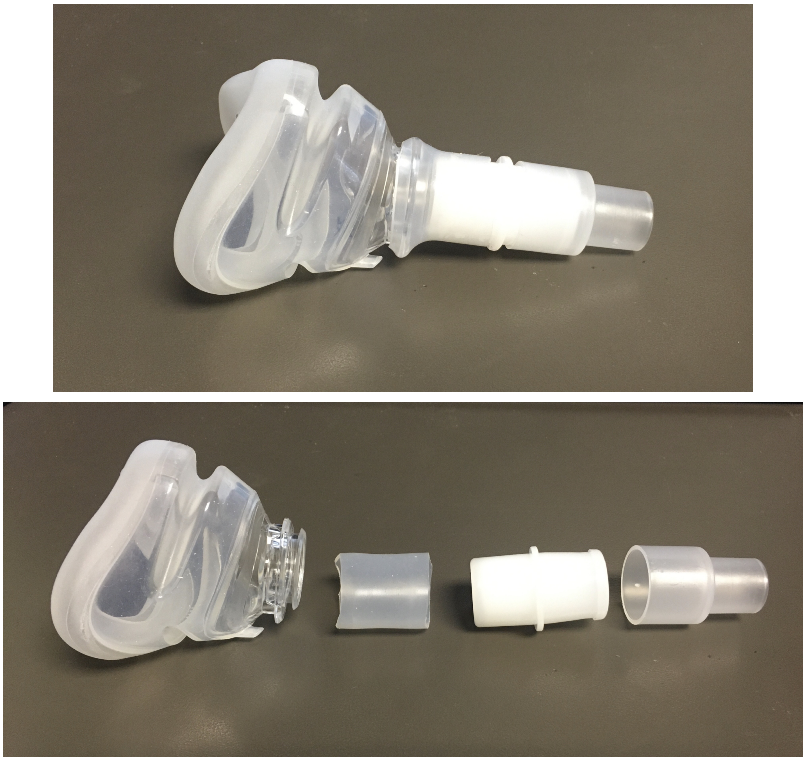

**Figure S7.** Odorant delivery system components and assembly overview.

##### Assembly

8. Cut a ~3.5 cm segment of silicone tubing (7/8" × 15/16").
9. Stretch the tubing over the front opening of the nasal mask to connect it to the one-way valve.
10. Insert the plastic adapter (supplied with the one-way valve) with the larger end over the one-way valve and the smaller end over the bag's open/close valve (Figure S8).

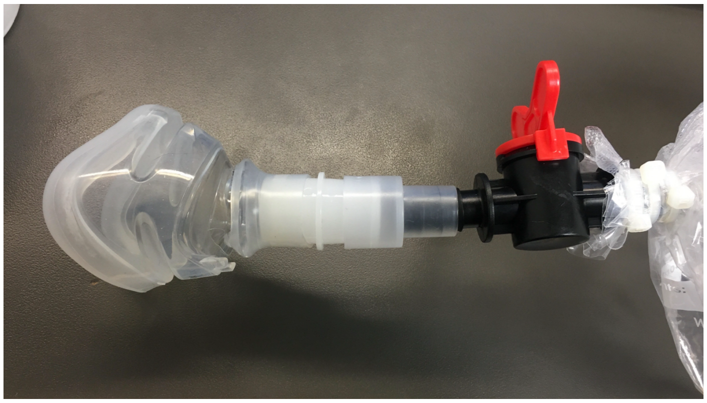

**Figure S8.** Connection detail showing the plastic adapter linking the one-way valve to the bag's open/close valve.

11. Mount the mask in the headgear frame and secure with the adjustable straps.
12. Fit the assembly to the participant, adjusting straps to ensure the mask creates a complete seal against the face.

    ***Note:*** A proper seal is critical to prevent dilution of the odorant stimulus with ambient air.

**Video Reference**

A video demonstration of the bag assembly technique is available at:
https://youtu.be/w_b2xQFffrY?t=69
